# A cobB like protein in *Oryza sativa indica* regulates the mitochondrial machinery under stress conditions

**DOI:** 10.1101/2022.04.17.488584

**Authors:** Sonali Khan, Nilabhra Mitra, Sanghamitra Dey

## Abstract

Sirtuins are ubiquitous in nature and are known to play an important role as metabolic regulators. In plants, these NAD^+^ dependent deacetylases are not well characterized. In this study, we are reporting a new member of sirtuin in *Oryza sativa indica*. It shares approx. 89% sequence identity with bacterial sirtuin, a class III sirtuin member. This protein is mostly present in mitochondria with trace amounts in the nucleus. It can physically interact with histones H3 and H4 and can specifically deacetylate histone H3 at K^9^ and K^18^ positions. In mitochondria, acetyl coA synthetase (AcS) and isocitrate dehydrogenase (IDH2) are the targets for its deacetylation. This removal of acetyl group is the mode of regulation under certain stress conditions. Thus, this is the first mitochondrial cobB targeting important plant machinery under changing growth environment. The OscobB deacetylase activity is not majorly affected by its products, NAM & ADP ribose but are sensitive to certain metal ions like Fe^2+^ and Mg^2+^. In contrast to some class III members, it does not have any ADP ribosyl transferase activity. In response to abiotic stress conditions like dehydration and low temperature, this enzyme can also mobilize to the nucleus to regulate the plant metabolism.

**Highlights:**

- A new class III member of the sirtuin family found in *Oryza sativa indica*.
- Under normal conditions, this protein is localized mostly in mitochondria, with trace amounts in the nucleus.
- This enzyme is capable of using both NAD^+^ and NADP^+^ as a substrate for catalysis.
- It specifically deacetylates the nuclear histone H3 at K^9^ and K^18^ positions.
- Mitochondrial proteins acetyl coA synthetase (AcS) and isocitrate dehydrogenase (IDH2) are the regulatory targets of OscobB.
- Under certain stress conditions in plants like dehydration, pathogenesis and low temperature, there is localisation of OscobB from mitochondria to nucleus.

## (1) INTRODUCTION

Plant proteins undergo post translational modifications (PTM) in response to environmental changes. PTM provides a wide range of protein diversity in this complex plant system to sustain changes in the surrounding environment. The process of acetylation/deacetylation are one of such modifications. Unravelling the mysteries of the PTM’s involvement in these complex mechanisms is highly intriguing.

The class III HDAC member, sirtuins are further classified into 5 classes (I-IV and U), depending on their amino acid sequence in the core domains as well as cellular localisation in both prokaryotes and eukaryotes [1]. Among mammalian sirtuins, SIRT1, 2, and 3 are class I sirtuins, having high homology to the yeast Sir2 and proteins such as Hst1, and Hst2, and exhibit robust deacetylase activity [1]. Class II sirtuins, including mammalian SIRT4, have no detectable deacetylase activity and instead show weak ADP ribosyltransferase activity [1,2]. Class III sirtuins, SIRT5 have weak deacetylase activity with several other catalytic abilities [3]. Most of the bacterial sirtuins also belong to this class. Few of the examples are Af1 from *Archaeoglobus fulgidus* (cobB1) [4], cobB from *Escherichia coli* [5] and *Salmonella enterica* [6], Pfsirt2A from *Plasmodium falciparum* [7]. Class IV sirtuins have both ADP ribosyltransferase and deacetylase activity (SIRT6 and SIRT7) [8,9]. Finally, class U sirtuins are found in several firmicute (gram positive) bacteria and *Thermotoga maritima* possessing sequence motifs that are intermediate between class I and IV, and have thus far only been observed in bacteria [1]. Collectively, the sirtuins play important roles in regulating genomic stability, energy metabolism, and stress resistance in a variety of organisms [10]. SIRT1, 6 and 7 are nuclear proteins in their localization and it is strongly evident that SIRT1 and 6 are linked to stress responses and aging processes [8,11,12]. On the other hand, SIRT3, 4 and 5 are mitochondrial proteins involved in aging processes and regulation of energy metabolism [1–3,13–16].

In comparison, plant sirtuins have not been studied in great detail. Two genes coding for sirtuins were identified in *Vitis vinifera*, namely VvSRT1, localized in the nucleus and VvSRT2, localized in both mitochondria and chloroplast [17,18]. VvSRT1 is noted to play a role in plant development whereas VvSRT2 is involved in maintaining the chromatin structure and regulation of gene transcription, indirectly linked to plant photosynthesis as well as leaf senescence. Two sirtuins genes have also been identified in *Arabidopsis thaliana*, namely *AtSRT1* and *AtSRT2*, both of which are localized in the mitochondria, although traces of AtSRT2 has been detected in nucleus as well [19–22]. AtSRT1 plays a role in auxin signaling [22] whereas AtSRT2 is a negative regulator of plant defense system by deacetylating PAD4 (phytoalexin deficient 4), EDSS (enhanced disease susceptibility) and SID2 (salicylic acid induction deficient) [22]. In tomato (*Solanum lycopersicum*), SoSRT1 is localized in the nucleus and SoSRT2 was seen to be localized in the nucleus as well as the cytoplasm [23]. These sirtuins were majorly involved in protein succinylation in tomato which regulated a number of physiological processes including photosynthesis. Glycine max has four sirtuins, GmSRT1, GmSRT2, GmSRT3 and GmSRT4 whose biological role has not been clearly studied [24].

Eukaryotes are known to possess several sirtuins for their cellular processes whereas bacterial cells mostly contain few of them, at the most, two [25]. In *E.coli*, which contains only one kind of sirtuin, CobB regulates chemotaxis by deacetylating CheY [26]. In bacteria these proteins are mostly present in cytoplasm. *E. coli* topoisomerase I (TopA) is also known to be regulated by N-Lysine acetylation. CobB regulates the negative DNA supercoiling activity by protecting the TopA from inactivation by non-enzymatic acetylation [27]. The strains of the enterobacter, *S.enterica* lacking this sirtuin, are unable to grow on propionate and in low acetate concentrations as the acyl-CoA synthetases (AcS) responsible for converting free acids into acyl-CoA derivatives are inactive [28]. *In vitro* NAD^+^-dependent deacetylase activity has also been described for numerous other class III sirtuins, like archeabacterial SIR2-Af1 (*Archaeoglobus fulgidus*) [4], *plasmodium falciparum* sir2a [7].

This study emphasizes on one of the sirtuins, found in *Oryza sativa var indica* (rice). Based on sequence analysis from Ensembl plants (https://plants.ensembl.org/index.html), there are three different sirtuins listed in *Oryza sativa indica*. OsSRT1 is closely related to the mammalian class IV member, SIRT6 [29] whereas OsSRT2 is related to SIRT4 [19]. OsSRT1 is required for the safeguard against genome instability and cell damage in rice plant [29]. This nuclear sirtuin functions as a regulator to repress glycolysis in seedlings and glycolytic genes expressions as well as inhibits transcriptional activity of glyceraldehyde-3-phosphate dehydrogenase (GAPDH) that is enriched on glycolytic genes promoters and stimulates their expression [30]. During oxidative stress, OsSRT1 reduces GAPDH lysine acetylation and reduces its accumulation in nucleus. In most of the cases, the deacetylation by OsSRT1 has been responsible for repression of the gene expression in different metabolic pathways. No relevant experimental data on OsSRT2 is available to portray its function in plants. We find that the third NAD^+^ dependent deacylase (UniProtID -A2XBC4_ORYSI) has 89% sequence identity with that of the bacterial protein, *Escherichia coli* cobB. There is no information available in literature regarding this third member in plants.

The molecular mechanism and kinetics of this class of protein need to be explored in plants. In this study, we have characterized OscobB protein and its catalytic abilities, found its localization in plants as well as discovering few of its interacting partners. This has provided us clues regarding OscobB’s role in mitochondria and regulation under stress conditions. The modulation of this enzyme’s actions is also studied here.

## (2) MATERIALS AND METHODS

All the reagents used in this study were purchased from HiMedia laboratory, India, Sisco Research Laboratories (SRL) India and Sigma-Aldrich, USA. Molecular biology related reagents: restriction enzymes, Taq DNA polymerase and T4 DNA ligase were from New England Biolabs, USA. Primers were synthesized from IDT DNA, USA. Recombinant histones (H1(M2501), H2A (M2502), H2B (M2505), H3 (M2503S), H4 (M2504S)) were purchased from New England Biolabs (NEB), USA. The histone H3Ac antibody sampler kit (9927T) and acetylated Histone H4 antibody sampler kit (ab218056) used were purchased from Cell Signaling Technology (CST) and abcam (USA), respectively. Antibody for AcS2 (ab66038), IDH2 (PHY0040S) and acetyl Lys (9681S) were purchased. H3K9Ac peptide (ARTKQTARK (ac)STGG), AcS2 peptide (KTRSGK(ac)IMRRI) were synthetically prepared from Bio Basic Inc, Canada. Custom made antibody for full length untagged OscobB was prepared from rabbit (Biobharati Lifesciences Pvt. Ltd, India).

### 2.1 Rice plant growth conditions

IR64 and Swarna varieties of rice seeds were first surface sterilized and laid on cotton sheets for germination in the dark for 48 hours. Then it is transferred to plant culture bottles with hydrophonics solution (MS media) and left at 25°C with 10h light and 14h dark cycles for about 3-4 weeks till 3 leaf stage. Cold or pathogen resistant varieties of rice seeds were also grown in similar way. For other stress conditions, three-week-old rice seedlings growing in MS medium were transferred to the medium supplemented with 500 mM NaCl. For exposing plants to dehydration stress conditions, the 3 week old plants were taken out of MS media for 2 days. All the plant samples were flash frozen and stored at −80°C.

### 2.2 Cloning of the *OscobB* gene

The RNA was isolated from the rice leaves using kit (Himedia-HiPurA™ Plant and Fungal RNA Miniprep Purification Kit) and cDNA was prepared using Invitrogen verso kit. The OscobB portion (269-501) of the OsI_09569 gene was amplified from the above cDNA using gene specific primers and was inserted into a modified pET28a vector between *NcoI* and *BamHI* restriction sites. This gene was also inserted in the pETM30 vector between *NcoI* and *BamHI* restriction sites to generate the GST-tagged construct. The sequences of the clones were further checked using DNA sequencing service.

### 2.3 Overexpression and purification of OscobB constructs

OscobB sequence containing an N-terminal 6xHis-GST tag in pETM30 vector and in modified pET28a vector (untagged protein) were expressed in *E.coli* BL21 (DE3) in LB medium. Cells were grown to A_600_ of 0.8 at 37°C and cobB expression was induced by adding 1mM isopropyl-1-thio-D-galactopyranoside (IPTG). The culture was further grown overnight at 18 °C. The cells were harvested by centrifugation at 4,000 x g for 30 mins and cell pellet stored in −20°C. Cells were resuspended in lysis buffer (50 mM phosphate buffer pH 7.5, 300mM NaCl, 10% glycerol, 1mM PMSF and 1mM DTT) and sonicated on ice. The cell debris were separated by centrifugation (14,000×g, 4 °C, 35 minutes). The supernatant of His-GST-OscobB was loaded on a Ni-NTA pre-equilibrated column with a lysis buffer. The column was washed thoroughly and protein was then eluted using an elution buffer (50mM phosphate buffer pH 7.5, 300mM NaCl, 10% glycerol, 1mM PMSF and 1mM DTT, 200mM imidazole). The supernatant of untagged OscobB was loaded onto an ion-exchange column (Bio-scale Mini Macro-Prep High Q Cartridge (BIORAD #732-4122) and the protein was eluted by using a NaCl gradient. Both the proteins were then desalted using HiTrap Desalting column (Sigma-Aldrich cat # GE17-1408-01) equilibrated with 50mM phosphate buffer pH 7.5, 100mM NaCl, 10% glycerol, 5mM PMSF and 5mM DTT. Both the purified proteins were resolved in 12% SDS-PAGE gel and stained using coomassie brilliant blue, found to be approx. 85% pure. The yield of the protein per litre of culture is approx. 1mg. All the pull down experiments were carried out using HIS-GST-OscobB protein. All the other analyses like the kinetic studies were carried out using the untagged OscobB protein.

The purified protein from untagged OscobB was further analysed by a size exclusion chromatography (SEC) passing 0.5mg protein through the SUPERDEX 200 (10/300 GL) column (GE Healthcare, USA) attached to AKTA pure 25 at the rate of 1ml/min. Buffer contained 20 mM phosphate buffer pH 7.5, 100mM NaCl, 10% glycerol, 5mM PMSF and 5mM DTT. The presence of OscobB was detected using absorbance at 280nm and SDS-PAGE gel.

### 2.4 Interaction studies of OscobB with histones: Ni pull down assay

Interaction experiments were performed by incubating 2μg His- GST- OscobB protein with 20μl pre-equiliberated Ni-NTA beads at 4°C for 2hrs. Then 2μg of recombinant histones (H3, H4, H1, H2A and H2B) were added to the mix in separate experiments and incubated for 2 hour at 4°C. Beads were centrifuged at 4,000×g, 4°C, 5 minutes to collect the flowthrough. The beads were further washed with buffer (50 mM phosphate buffer pH 7.5, 300mM NaCl, 10% glycerol, 5mM PMSF and 5mM DTT) several times. Finally, the beads were mixed with sample dye and boiled. The proteins stuck in the beads were then resolved using 12% SDS-PAGE gel and stained using coomassie brilliant blue dye. Similar experiments were carried out to analyse the interaction of OscobB with AcS, IDH2 by mixing the beads with mitochondrial extracts prepared using cell fractionation kit (ab109719).

### 2.5 *In vitro* lysine deacetylation activity assay

For Western blot analysis, the reaction was performed at 37°C for 2 hrs. Buffer prepared was 50mM Tris pH 7.5, 150mM NaCl, 5mM DTT and 1mM NAD^+^. A 40μl of reaction mixture was prepared containing 2μg untagged OscobB, 30μg histone plant extract and final volume was made up by buffer. The reaction was stopped by adding sample dye and boiling for 5mins. The mix was then resolved using 12% SDS-PAGE gel and the protein bands on the gel were transferred on nitrocellulose membrane. Immunoblot analysis was done by initially blocking the membrane in 5% skimmed milk + TBS for 20mins. Rabbit monoclonal acetyl-lysine primary antibody (1:1000) followed by an HRP-conjugated Goat Anti-rabbit secondary antibody (1:5000) was used to detect the deacetylation activity of the protein. Time point experiment was performed by collecting the reaction mix after every 15 mins, mixed with sample dye and boiled. Further western blot analysis was performed using antibodies specific for histone H3 lysine sites (9927T) as well as for some histone H4 lysine sites (8346T). Further, deacetylation experiments were performed keeping the concentration of OscobB protein as constant and gradually increasing the concentration of NAD^+^ from 100 μM to 800 μM. A reaction mix was made up of untagged OscobB protein (0.8μM), nuclear protein with histones extracted from rice leaves (40 μg) and buffer (50mM Tris pH 7.5, 150mM NaCl, 5mM DTT). It was then incubated at 25°c for 2 hours followed by blotting on a nitrocellulose membrane. The blot was blocked using 5% skimmed milk + TBS following which western blot analysis was performed using rabbit monoclonal acetyl-histone antibody (1:1000) followed by a HRP-conjugated Goat Anti-rabbit secondary antibody (1:5000). The experiment is performed in triplicates. Detection of the blot was done using the Biorad ChemiDoc MP imaging system and quantification was done using imageJ. Similar dot blot experiments were performed using NAD^+^ analogs in place of NAD^+^ in the reaction mix to determine their involvement in the deacetylation reaction. Kinetic parameters for histones, AcS and IDH2 deacetylation were carried out using the similar dot blot method. Saturation kinetics using Non-Linear regression (Michaelis Menten or *k_cat_* Model) was prepared using GraphPad Prism version 9.1.2 (San Diego, USA).

### 2.6 Intrinsic tryptophan fluorescence binding studies

Experiments were performed at room temperature where untagged OscobB protein (0.8 μM) was incubated with NAD^+^ (0 to 600μM) and the emission spectrum was monitored by exciting the samples at 295nm. The resultant emission was recorded from 305 to 480 nm using Hitachi Fluorimeter with a slit window of 2.5mm. Binding studies of OscobB with other ligands were also carried out in a similar manner in Biotek SYNERGY H1 microplate reader. The Kd values were calculated using one site binding equation in Graphpad prism version 8.3.0 (San Diego, USA).

### 2.7 ADP ribosylation assay of OscobB

The ADP ribosylation reaction was performed at 25°C for 1 hr. Buffer composition for the reaction was 50 mM Tris pH 7.5, 150 mM NaCl, 10 mM DTT and 75 μM NAD^+^ and 40 μM 6-biotin-17-NAD (Bio-NAD^+^, R&D, Minneapolis, USA). The reaction mix containing 0.8μg recombinant OscobB as well as OscobB with whole protein were then resolved using 12% SDS-PAGE gel and transferred on a nitrocellulose membrane. The immunoblot analysis was done by initially blocking the nitrocellulose membrane in 5% skimmed milk in TBS for 30 mins. Rabbit monoclonal anti-biotin antibody (1:5000) was used to detect the biotinylated ADP ribose moiety of the protein. This was followed by incubation with HRP conjugated secondary antibody (1:10000) and developed with ECL solution (BioRad).

## 3. RESULTS AND DISCUSSION

### 3.1 Structural bioinformatic analysis of OscobB

While studying different NAD^+^ dependent deacetylases in *Oryza sativa indica* we came across this new protein sequence (OsI_09569) in the Ensembl plants database (http://plants.ensembl.org). Blastp search of this putative sirtuin protein sequence finds it to be a member of class III sirtuin family. It has sequence identity of approx. 89% with *E.coli* cobB and 37% with human SIRT5 (class III members). Among plant sirtuins, OscobB shares approx. 23% and 29% sequence identity with OsSRT1 and OsSRT2, respectively. **(Fig1A)** In its amino acid sequence, we could find the signature motifs of sirtuin, GAGISAESGIRTFR and YTQNID. Using the three-dimensional structure of *Ecoli* cobB (1S5P) as a template, a homology model of OscobB was prepared using the SWISS-MODEL webserver [31]. This structure clearly delineates the small zinc binding domain from the major active site containing the Rossmann fold subunit **(Fig1B)**. The presence of the Rossmann fold suggests a binding site for NAD^+^/NADP^+^ in this protein. The zinc ion is pointed away from the active site, suggesting its non-participation in the direct catalytic activity of sirtuins. But presumably, it plays an essential role in maintaining the structural stability of the catalytic core domain. The sirtuin structure is known to partially collapse in the absence of the Zn^2+^ ion [32]. In most sirtuins, four cysteine residues are involved in coordination with the Zn^2+^ ion in a tetrahedral geometry to maintain the structural integrity of the sirtuin. Here the cobB structure follows the zinc binding motif (Cys-X_2_-Ser-X_15_-Cys-X_2_-Cys), with the Zn^2+^ ion tetrahedrally coordinated between three Cys and one Ser residue (Cys^118^, Ser^121^, Cys^137^ and Cys^140^). Interestingly, in the *E. coli* cobB structure [33] it is noticed that the Ser is not involved in the coordination with Zn^2+^ rather a nearby Cys plays the role. This Cys residue is also conserved in OscobB (Cys^139^). The two domains in this structure are linked by several variable loop regions. The conserved catalytic core is present in area in the form of a tunnel **(Fig1B)**. Overlay of OscobB model with HsSIRT5 and PfSir2 (other classIII members) showed a good superimposition in Rossmann fold region, much variation is noticed in the Zn binding domain. This provides a variability in substrate binding at the active sites of the said sirtuin members **(Fig S1)**.

**Fig 1.**
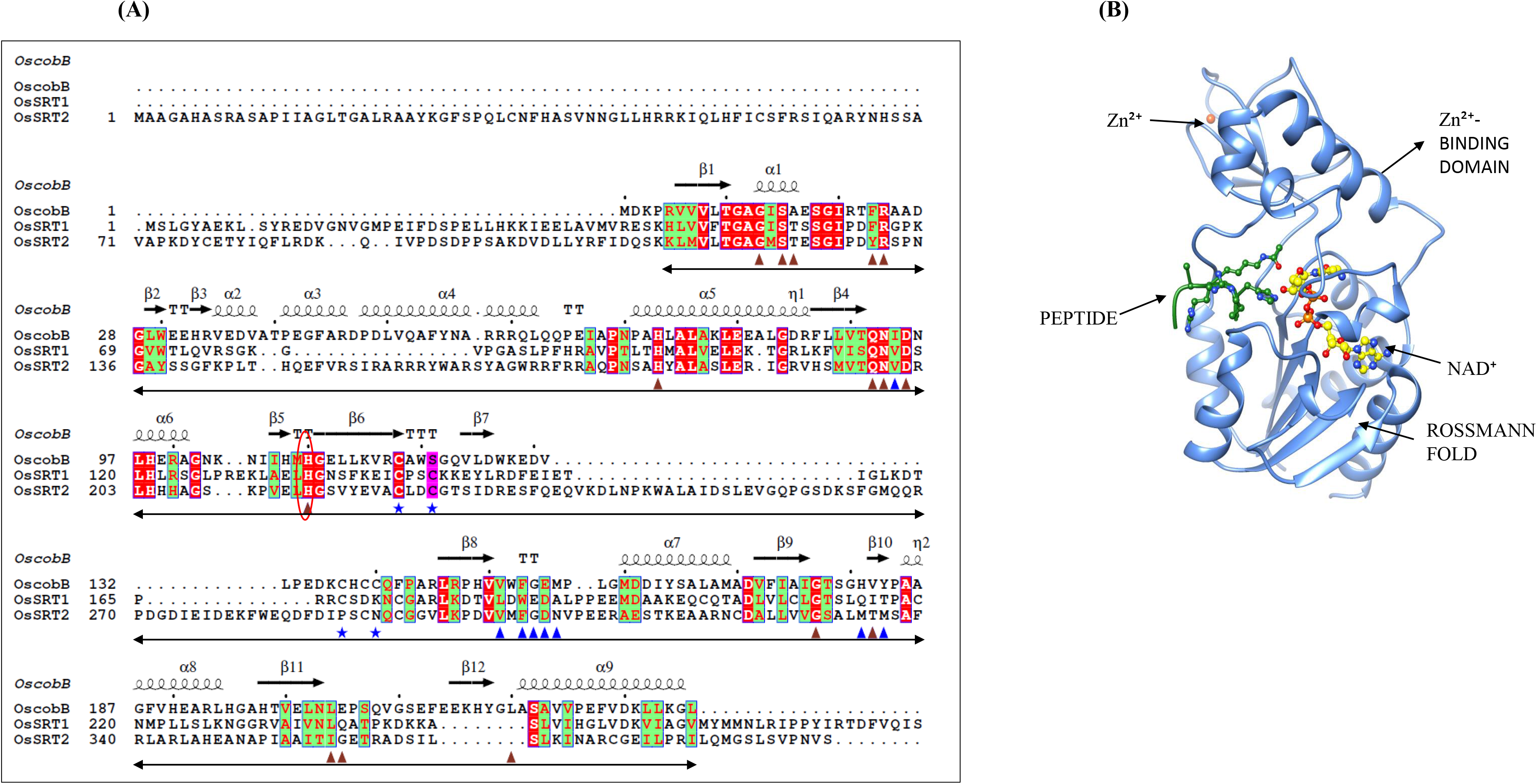
Structural analysis of OscobB: **(A) Sequence analysis of all the rice sirtuins:** Multi sequence alignment of rice OscobB (Uniprot ID A2XBC4) with OsSRT1 (Uniprot ID B8ARK7) and OsSRT2 (Uniprot ID B8BNG4) is shown here. The residues which line the NAD^+^ binding region are shown in maroon triangles, peptide binding residues with blue triangles and cysteines involved in zinc binding are shown with blue stars. The catalytic base, His^110^ is shown with a red circle. The underlying black arrows demarcate the catalytic core region of the sirtuins. The sequence of OsSRT1 in the alignment is also truncated for clarity. The sequences are aligned using clustalW and displayed using ESPript 3.0. **(B) Cartoon representation of OscobB structure:** The three dimensional model of OscobB is prepared by using Swiss-model webserver, with *E.coli* cobB (PDB ID-1S5P) as the template. The sirtuin structure is bound to H3K9Ac peptide (green ball and stick) and NAD^+^ (yellow ball and stick representation). The OscobB structure is divided into the Rossmann fold region and a variable zinc-binding domain. The NAD^+^- and substrate binding regions are shown to occupy different areas on the enzyme.

Eukaryotic sirtuins have amino (N)– and carboxy (C)–terminal extensions that are divergent among the members of this family. Its variable terminal regions are responsible for oligomerisation, binding to accessory proteins or substrates, and its regulation. These are also required for enzyme catalysis as well as its diversity. In OsSRT1, the extra long C-terminus is required for substrate binding and its catalysis [34]. In comparison, OscobB do not have any extended N- and C-terminal regions as can be seen from the sequence alignments **(Fig1A)**. It is rather a compact small protein.

### 3.2 Subcellular localisation and tissue expression

The OscobB expression analysis in 3 week old rice plants suggests that this protein is present mostly in stem and leaves **(Fig2A)**. Under normal conditions, after extraction of different organelles of rice leaves, western blot analysis of these fractions using OscobB specific primary antibody was carried out. It showed the presence of this enzyme mostly, in mitochondrial fractions and to a lesser extent in the nucleus (21%) **(Fig2B)**. TargetP and Mitofate analysis of the protein sequence of OscobB also suggested its mitochondrial localization [35–37]. This is the first instance that the cobB protein is present in two different organelles. In comparison, literature studies on other rice sirtuins report OsSRT1 to be mostly localized in the nucleus [38,39] and OsSRT2 to be a mitochondrial protein [39]. No data is yet available for interorganellar localization in these plant sirtuins. Our studies predict that this enzyme OscobB might be having inter organelle functions in rice plants. It would be interesting to find out under what cellular conditions, this protein localizes between the two organelles. Here, we can speculate that in response to certain stress conditions, OscobB can localise from mitochondria to nucleus to regulate the gene expression as seen in their human homolog SIRT5 [40]. Various groups have detected the localization of human SIRT5 from mitochondria to cytoplasm, peroxisome and nucleus [41,42]. Also, in human SIRT3, it is targeted from nucleus to mitochondria in response to stress conditions [43,44]. However, using different nuclear localization signal (NLS) prediction softwares, we were unable to locate the NLS sequence in OscobB protein. So, there is also a possibility of presence of a carrier or accessory protein, which might help this enzyme to move from mitochondria to nucleus under different cellular conditions.

**Fig 2.**
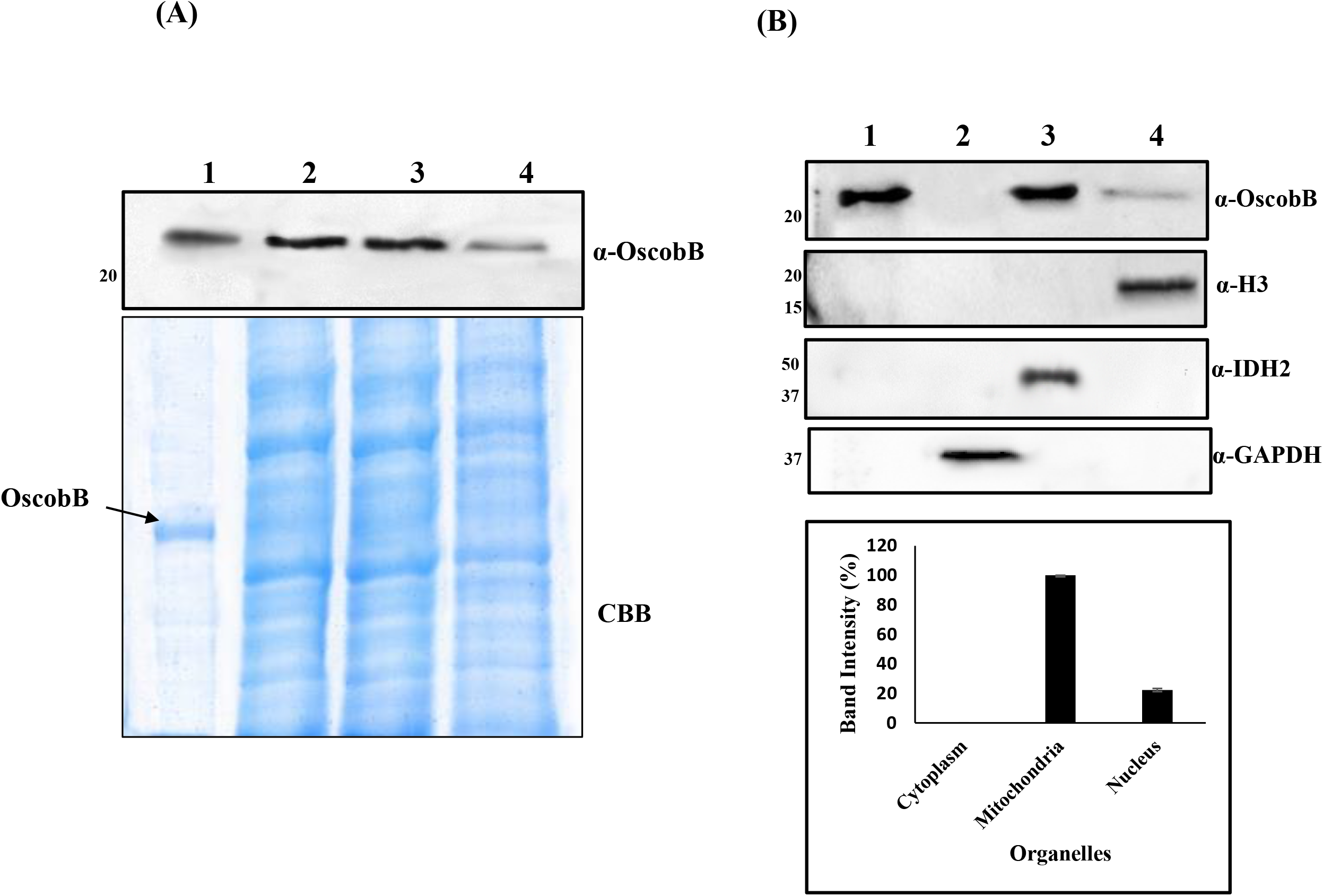
Localization of OscobB. (A) Tissue localization of OscobB: Equal amount of plant extract (7μg) is loaded on a 12% SDS-PAGE gel. Lane 1- purified recombinant OscobB, lane 2- leaf extract, lane 3- stem extract, lane 4- root extract. The presence of OscobB was confirmed by western blot analysis using OscobB-specific primary antibody. OscobB protein was observed to be equally dominantly present in stem as well as leaf extract. However, the root extract showed a lesser amount of this protein. (B) Analysis of the subcellular OscobB expression in leaf under unstressed condition. Equal amounts of cell-organellar extract (approx. 10μg) were loaded on a 12% SDS-PAGE gel. Lane 1- purified OscobB, lane 2- cytosolic extract, lane 3- mitochondrial extract, lane 4- nuclear extract. Nuclear, mitochondrial and cytoplasmic extracts were verified using specific marker antibodies, histone H3, IDH2 and GAPDH respectively. OscobB was seen to be predominantly present in the mitochondria, followed by the nucleus. The bar graph depicts the amount of OscobB protein in plant extracts based on its western blot band intensity calculated using ImageJ software.

### 3.3 NAD^+^ dependent deacetylase activity of OscobB and its possible substrates

To further characterize the catalytic abilities of recombinant OscobB as an enzyme, western blot experiments were carried out. As mentioned earlier, the untagged protein was purified using ion exchange chromatography using NaCl gradient buffer. Analytical SEC analysis of the untagged OscobB protein showed a monomeric protein (25kDa) in phosphate buffer solution at pH 7.5 **(Fig S2)**.

We were further interested in finding its physiological substrates in plants. Due to its nuclear localisation, the Ni pull down experiments of OscobB with histones were performed. The experiments showed the binding of the histones H3 and H4 with this enzyme **(Fig3A)**. However, the interaction studies of OscobB with the other histones H2A, H2B and H1 showed no binding with these proteins **(Fig S3**). It is quite possible that the histones H3 and H4 are the probable substrates for OscobB action in nucleus. A time point experiment for the deacetylase activity was performed using the anti-acetyl western blot. In this reaction, OscobB was mixed with NAD^+^ and leaf nuclear extracts. The enzyme was able to completely deacetylate the histone H3 (1μg) within 2 hour time (**Fig3B)**. So, we can say that the purified OscobB enzyme is active and shows NAD^+^ dependent deacetylase activity. It is well known that sirtuins can remove the acetyl group present on the lysine residues of the substrate. Using histone site specific western blot analysis, we detected the deacetylation at K^9^ and K^18^ positions of H3 by OscobB. Even though this protein interacted with histone H4, it did not deacetylate this target **(Fig3C and S4)**. There is a possibility that H4 may act as a substrate for other possible catalytic actions of OscobB. Further, western blot analyses were conducted to calculate the kinetic parameters of these H3 deacetylations. The *K*_m_ value for NAD^+^ and H3K9Ac peptide for OscobB were found to be 139 ± 23 μM and 104.7 ± 6 μM, respectively **(Fig3D)**. The reaction rate for the H3K18Ac deacetylation is weaker compared to H3K9Ac with a *k_cat_/K_m_* of 1.3 X 10^6^ M^−1^s^−1^. In plants, nuclear OsSRT1 also deacetylates H3K9Ac linking it to DNA repair mechanisms under oxidative stress conditions [38]. It needs to be seen how the deacetylation of the same residue by OscobB is related to its functional mechanism in plants.

**Fig 3.**
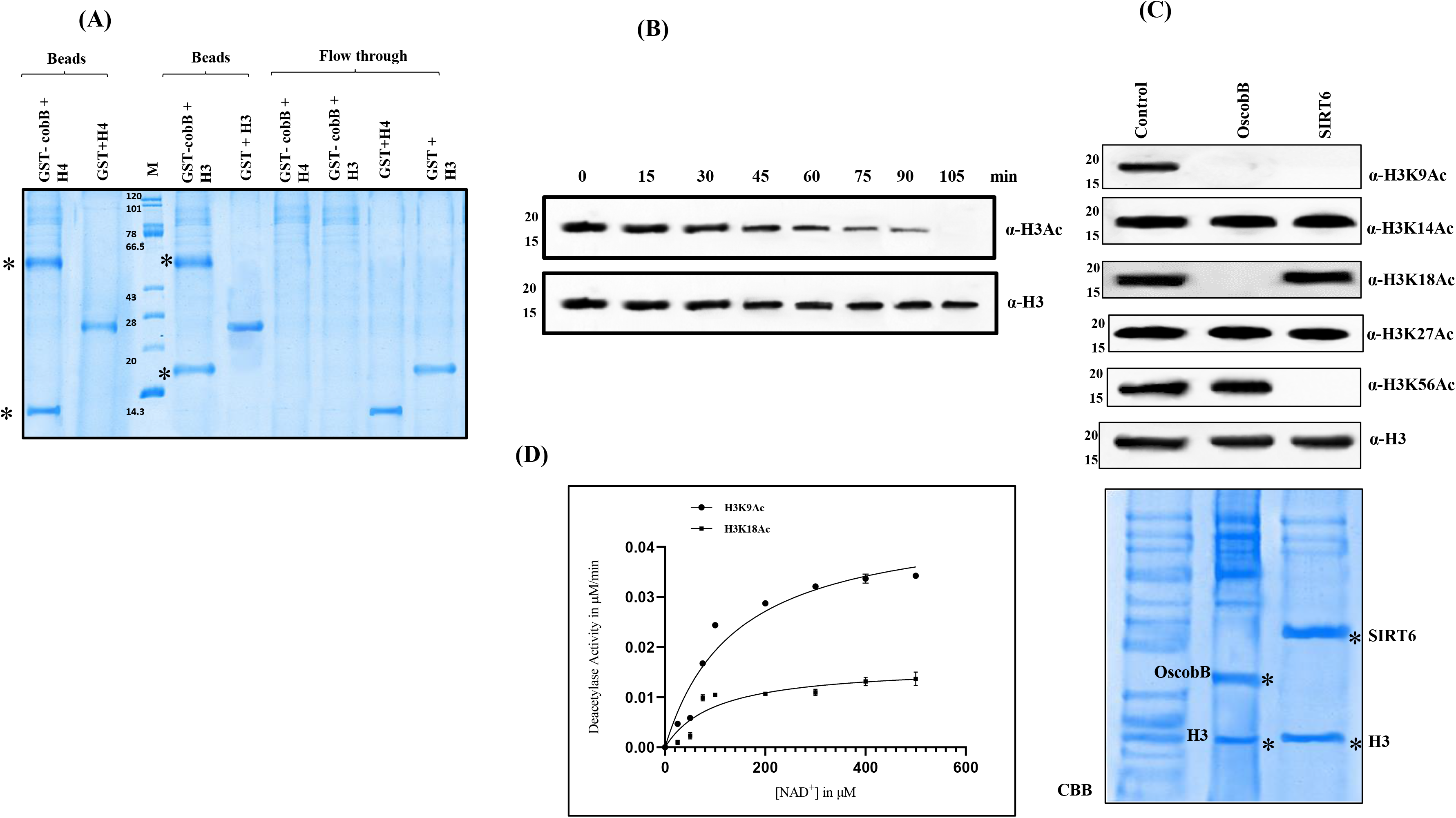
OscobB- histone binding and deacetylation activity. (A) GST pull down assay was performed to detect the binding of HIS-GST-tagged OscobB with recombinant histones H3 and H4 (NEB). Here, 12% SDS-PAGE gel shows OscobB can physically bind to both the histones H3 and H4. GST protein alone was used as a negative control in lanes 8 and 9 suggesting that the histones are not non-specifically binding with GST. (B) Detection of deacetylase activity of OscobB: Deacetylation assay of histone H3 from rice leaf nuclear extract was performed at 28°C. The reaction mixture containing OscobB was collected at different time points (0-105 mins). Sample reaction was stopped by boiling them with sample dye and then loading on a 12% SDS-PAGE gel. Western blot analysis was performed using H3Ac primary antibody to detect the extent of deacetylation by OscobB. Here the loading control is histone H3. (C) Detection of the deacetylation of specific lysine residues in histone H3 by OscobB. Lane 1- Empty pET28a vector, lane 2- purified recombinant OscobB, lane 3- purified HsSIRT6. Anti-H3 antibody was used as loading control for all the western blots. Empty vector and HsSIRT6 were used as negative and positive control, respectively. Coomassie brilliant blue gel shows the reaction mix containing purified OscobB along with histone H3 in lane 2 and purified HsSIRT6 along with histone H3 in lane 3. (D) Saturation kinetics for the OscobB enzyme activity: MM plot showing the OscobB deacetylation of H3K9Ac and H3K18Ac from rice leaf nuclear extract using varied concentrations of NAD^+^ (0-600 μM). The non-linear regression plots were calculated using Graphpad Prism 8.3.0. The error bar depicts the S.D.; n=3. Further kinetic parameters (*K_m_* and *k_cat_* values) were calculated using these plots.

Additional Ni pull down experiments of OscobB with the mitochondrial extract could catch the interaction of this enzyme with acetyl coA synthetase (AcS) and isocitrate dehydrogenase (IDH2) **(Fig4A)**. The identity of the interacting partners was confirmed by AcS and IDH2 specific antibody. Acetyl coA synthetase plays a major role in fatty acid and polyketide biosynthesis and is helpful in plant seed development [45]. It catalyses the synthesis of acetyl coA using acetate, coA and ATP. OscobB could deacetylate AcS with a rate of 0.06 ± 0.006 μM/sec (**Fig4B)**. There are instances of human AcS getting deacetylated by SIRT3 [46]. Here, the lys^648^ of HsAcS gets deacetylated, which is also conserved in rice AcS (Lys^642^) and can be the probable site of deacetylation by OscobB. Further, a recent study on *Arabidopsis* AcS shows that its activity gets modulated by deacetylation at lys^622^ (approx. 500 fold) [47]. Though the authors did not report the deacetylase enzyme involved in this process. Deacetylation of AcS in *S. enterica* allows it to grow in acetate and propionate medium [28]. Perhaps, to combat certain stress conditions in plants, AcS activity also increases with the subsequent deacetylation by OscobB. Under certain environmental conditions like tolerance against the pathogen attack in rice plants, there is an increase in the expression of OsAcS. It will be really useful to unravel the mechanism behind this, as much of this information is not quite clear in plants.

**Fig 4.**
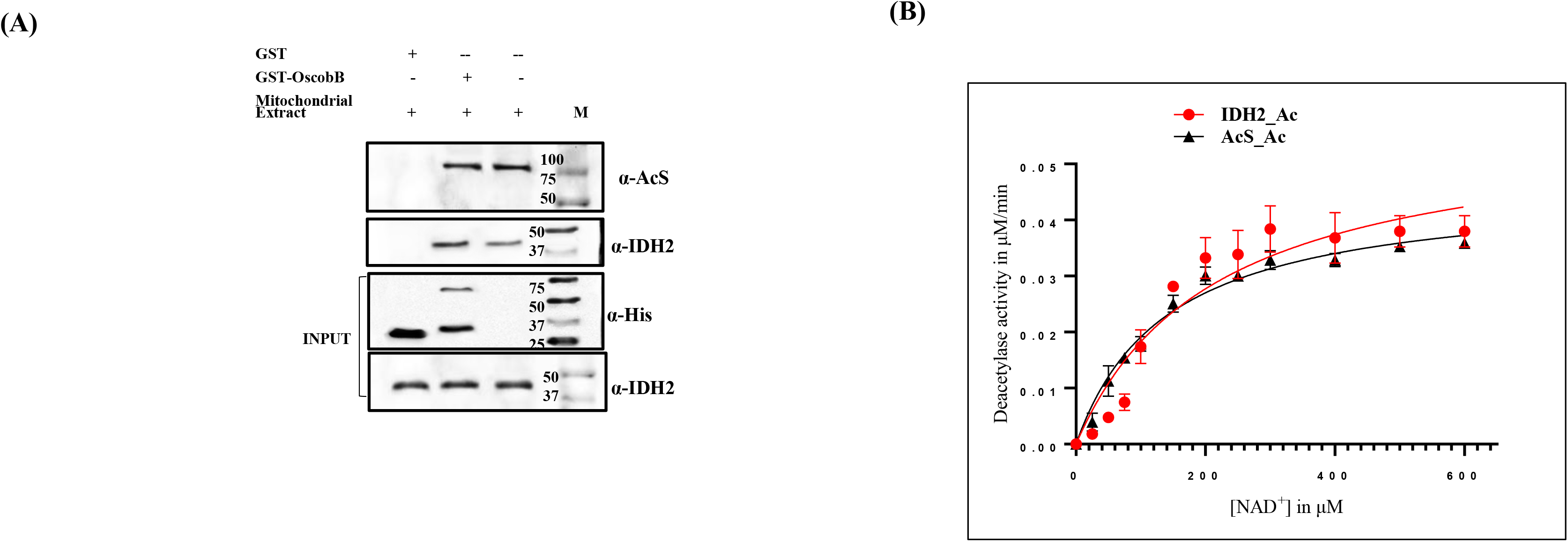
Non histone targets of OscobB deacetylation. (A) OscobB physically interacts with mitochondrial proteins AcS and IDH2. Ni pull down assay was performed to determine the interaction of His-GST-tagged OscobB to AcS and IDH2 extracted from leaf mitochondria. The resultant protein complexes were pulled down using Ni-NTA beads. Beads were mixed with sample dye and boiled and further resolved on a 12% SDS-PAGE gel. Western blot analysis was done using anti-AcS/IDH2 primary antibody to ascertain the identity of AcS. Input Mitochondrial extract was verified using specific organelle marker antibodies, IDH2 as well as anti-His antibody. (B) The MM plot showing the deacetylation kinetics of acetylated AcS peptide and IDH2 using varied concentrations of NAD^+^ (0-600 μM). The resultant plot was calculated using Graphpad Prism 9.3.0. The error bar depicts the S.D.; n=3. Further kinetic parameters (*K_m_* and *k_cat_* values) were calculated using these plots.

In rice mitochondria, the other protein under regulation is isocitrate dehydrogenase (IDH2). The site for deacetylation in IDH2 is conserved lys^391^ (equivalent to Lys^413^ in human IDH2). It is the site for regulation as deacetylation also increases the activity of human IDH2. Our Ni pull down results show the direct interaction of OscobB with IDH2 from mitochondrial extracts, with deacetylation rate of 0.055 ± 0.006 μM/min. So, we can speculate that OscobB can be the deacetylase involved in the regulation of IDH2 in mitochondria. IDH2 acts as an important control point in the TCA cycle in plants generating energy. There is no information available regarding the effect on plant IDH2 activity on its deacetylation. It is quite obvious that this enzyme is affected under certain stress conditions and its activity is regulated by deacetylation [48,49].

Sirtuins are very much dependent on NAD^+^ for their activities. Are any other analogs of NAD^+^ capable of performing the same task? Deacetylase activity of OscobB using NAD^+^ and its analogues showed a wide range of enzyme responses. This experiment showed that apart from NAD^+^, NADP^+^ and thio-NAD^+^ can also be used as cofactor for OscobB deacetylation, but with reduced efficiency. As expected, we did not see any deacetylase activity in the presence of NADH. We found comparable OscobB deacetylation using both NAD^+^ or NADP^+^ as substrates (**FigS5A &B)**.

There are no NAD^+^ bound structures available with the template sirtuins. To further understand the binding of NAD^+^ with OscobB, we have docked the NAD^+^ in the OscobB model using autodock vina (−9.7 kcal/mole) (**FigS5C)**.

### 3.4 Binding studies of NAD^+^ and its analogs: relation between its binding and deacetylase activity

It is known that sirtuin catalysed reactions are always bisubstrate. It involves hydrolysis of NAD^+^ coupled with protein deacetylation. In certain human sirtuins, NAD^+^ cannot bind to the active site in the absence of an acetylated peptide [50,51] whereas in human Sirt6 and 7, NAD^+^ can interact with the enzyme in the absence of acetylated peptide [52,53]. This mechanism of substrate binding at the active site is not known in any of the cobB proteins. So, to understand this molecular mechanism in the OscobB reaction, the intrinsic fluorescence experiments of this protein in presence of NAD^+^ and H3K9Ac peptide were performed. For this we have taken advantage of the intrinsic Trp residues (Trp^30^, Trp^99^ and Trp^152^) near the active site i.e. binding sites of the substrates. An emission peak at approx. 340nm was observed when the OscobB sample was excited at 295nm. On NAD^+^ binding, there was a decrease in fluorescence signal at 340nm, indicating tight binding with Kd of 30±2 μM (**FigS6A)**. Similar pattern was also observed on addition of thio-NAD^+^ and NADP^+^, indicating that all these analogues of NAD^+^ tend to bind at the active site of OscobB whereas in case of NADH, no change in fluorescence emission was noticed, indicating that NADH is not involved in binding to OscobB at its active site. So, this study also suggests that NAD^+^ can bind in absence of an acetylated peptide.

Next, the pattern of binding of H3K9Ac peptide with OscobB was followed. Addition of this peptide alone to the protein sample showed less deviation in the fluorescence intensity at 340nm on sample excitation at 295nm **(FigS6B)**. This might be the indication of weak binding to OscobB. However, we think that there is proper binding of the peptide in presence of NAD^+^ with the formation of ADPr, indicated by the decrease in the fluorescence signal. Thus, the H3K9Ac peptide alone is weakly binding with the enzyme and NAD^+^ is required for the second substrate binding in case of OscobB. **(FigS6C)**. Perhaps, NAD^+^ binding induces conformational change at the active site facilitating the binding of the second substrate.

### 3.5 Modulation of OscobB enzyme activity

Sirtuins are important regulatory enzymes in the biological system which can affect many different metabolic pathways in various organelles. That’s why it is necessary that they themselves are also controlled in some way in plants. There is limited or no knowledge regarding the regulation of sirtuins’ actions in plants. Product inhibitions are mostly one of the ways to regulate an enzyme. In the sirtuin bisubstrate reaction, along with deacetylated lysine, NAM and 2’O-acetyl ADP-ribose (OAADPr) are the two products that are released on NAD^+^ hydrolysis. In this case, the effect of varied concentrations of nicotinamide (NAM) on this enzyme reaction was determined. The IC_50_ value of NAM inhibition for OscobB deacetylation was found to be 253 ±45 μM. ADP-ribose, which is the analog of the other product of this chemical reaction (O’AADPr), also inhibited OscobB activity with IC_50_ value of 410 μM **(Fig5A)**. Similar insensitivity to NAM inhibition has been observed for the other class III member, human SIRT5 (0.7-1.6 mM) [54,55]. Trichostatin A which could inhibit class IV sirtuin (Hssirt6) had shown no effect on OscobB action, even at 1mM concentration. Further, we have looked at the effect of different metal ions like Cu^2+^, Ca^2+^, Mg^2+^, Mn^2+^ and Fe^2+^on the OscobB deacetylase activity. This reaction is mostly inhibited by Fe^2+^ and Mn^2+^ ions **(Fig5B)**. The mechanism of this inhibition is not quite clear but may indicate the effect of metal toxicity on plant growth. Natural polyphenol resveratrol, a well-known activator of sirtuins, increases OscobB deacetylase activity by 50 % at 51μM concentration (EC_50_) **(Fig5C)**.

**Fig 5.**
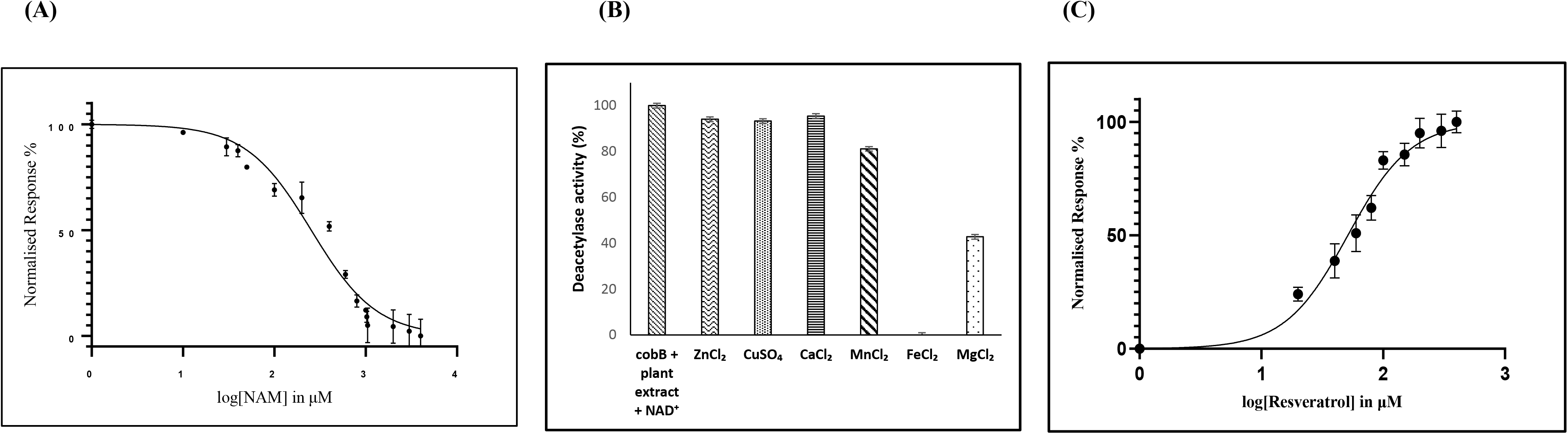
Modulation of OscobB deacetylase activity. (A) Dose-dependent effects of NAM on OscobB deacetylation activity using acetylated H3K9 as substrate. The reactions were carried out in triplicates by dot blot method using varied concentrations of nicotinamide (NAM) (0-1M). (B) Effect of metal ions on deacetylase activity of OscobB. The bar graph compares the effect of metal ion-dependent deacetylation of histone H3 from nuclear extract. 0.5mM concentration of each metal salts were used in the reaction and their effects were deciphered using western blot analysis. The error bar depicts the S.D.; n=3. (C) Dose-dependent effect of resveratrol on the deacetylase activity of OscobB. Histone H3 extracted from rice leaf nuclear extract was used as a substrate. The reactions were performed in triplicates using dot blot method using varied concentrations of resveratrol (0-400μM). Log (resveratrol) vs. response curve for the EC_50_ determination using Graphpad Prism. (n=3; error bar: s.d.)

## 4. CONCLUSIONS AND FUTURE DIRECTIONS

In this study, we have reported a cobB like protein in plants, which belongs to class III sirtuin family. In most of the organisms where cobB enzyme was studied, it is localized in cytosol. Here, in plants we find it to be majorly present in mitochondria. This enzyme is known to have multiple catalytic abilities. OscobB has good deacetylase activity but no ADP ribosyl transferase activity (**FigS7)**. This contrasts with the previous study on *S typhimurium LT2* cobB (89 % sequence identity) [56]. There is also a report on ADP-ribose transfer on BSA by *E.coli* cobB [16] and pfSIR2a [7], another class III sirtuin family member. In future we can explore other deacylase activities of OscobB in plants. The presence of the conserved active site residues (Tyr^56^ and Arg^59^) in this protein predicts its desuccinylase/ demalonylase activity as the active site can accommodate more acidic substrates [3]. It needs to be further seen what type of deacylase reaction is most favored by OscobB. Kinetically, this enzyme can utilise both the cofactors NAD^+^ and NADP^+^ for its deacetylase reaction. With this catalytic ability, it can specifically remove the acetyl group of K^9^ and K^18^ positions of nuclear H3. OscobB showed weak product inhibition. As we see that NAM did not have much effect on this enzyme, it will be worthwhile to look for its other regulatory mechanism. However, OscobB activity is greatly affected by metal toxicity. We can relate this to metal poisoning in soil which hampers the plant growth. Perhaps, OscobB activity can also get modulated by PTM of certain sites in its folds.

Sirtuins are seen to be an excellent player in response to metabolic stress. The class III sirtuin member, cobB is mostly prevalent and well-studied in prokaryotes. In *E.coli*, cobB has been a versatile player. We have highlighted some of its deacetylation targets in the rice plant. It needs to be seen what this enzyme can do in plants where it can localise in two different organelles. **(Fig6)**.

**Fig 6.**
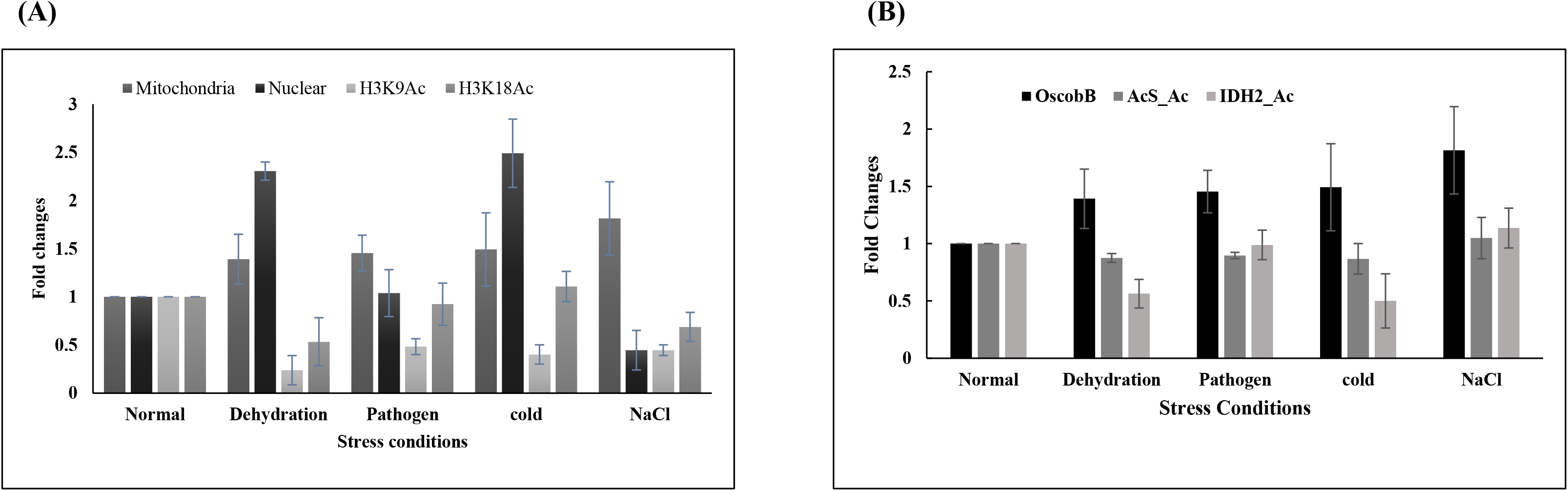
Relationship between the OscobB expression and varied factors under different stress conditions. (A) Western blot analysis of the cell extracts of rice leaves, exposed to varied stress conditions, was performed using OscobB and site-specific acetylated lysine antibodies of H3 to detect the intensity of their presence in those samples. The band intensity is measured using ImageJ and the values are normalized by an external control before plotting (Microsoft Excel). The histograms suggest a possibility of increase in the deacetylation of H3 sites (H3K9Ac and H3K18Ac) by the upregulated nuclear OscobB under dehydration and low temperature conditions. Here the profile for both the nuclear and mitochondrial OscobB under varied stress conditions are shown. (B) Western blot analysis of the mitochondrial extract of rice leaves exposed to varied stress conditions was performed using OscobB and acetyl-lysine antibody to detect the extent of acetylation in AcS and IDH2 proteins. Controls were also run to detect the presence of AcS and IDH2 proteins. The band intensity of the immunoblots were measured using ImageJ and the values were normalized by an external control before plotting (Microsoft Excel). The histograms suggest a relationship of increased deacetylation of these mitochondrial proteins with the upregulated OscobB under dehydration and low temperature conditions.

Under abiotic stress conditions like dehydration, cold and pathogenesis we found an increase in OscobB expression in nucleus than in mitochondria. Low temperature and dehydration cause a lot of damage to plant growth with production of oxidative stress in cells. This results in lots of ROS production and triggering the activation of certain enzymes to tolerate the situation [57]. It is possible that in response to these stresses, cobB remains in the nucleus from mitochondria to regulate certain pathways to adapt to these changes. As we are unable to locate the NLS or MTS in this protein sequence, it is possible that it requires accessory proteins for this localisation between mitochondria and nucleus. In relation to this, we also observe that the deacetylation of H3K9Ac and H3K18Ac in the nucleus increases under cold and dehydration. There is also increased expression of AcS under certain stress conditions. As we know, metabolic AcS are mostly regulated by acetylation. In this case, OscobB can act as a regulatory deacetylase in mitochondria. Thus, OscobB can regulate the level of acetyl coA by deacetylating AcS. It is the first reported mitochondrial protein to be deacetylated by sirtuins in plants. It is known that acetylation of AcS inactivates the enzyme. Under certain stress conditions, OscobB deacetylates AcS, which in turn gets activated. Earlier it was not known that the AcS modulation in plants can be due to OscobB action. Now we can say that the sirtuin mediated regulation of AcS activity by its deacetylation is well conserved from bacteria to humans to plants.

There is also not much information available regarding the plant IDH2 activity and its regulation in mitochondria. This study sheds some light on this aspect of such an important protein. Stress may affect the mitochondrial oxidation leading to more energy generation in plants. Deacetylation leads to activation of this enzyme. Further investigation needs to be done to understand the role of OscobB in this regard.

## SUPPORTING INFORMATION

**Figures S1–S7**

## ACKNOWLEDGEMENT

This work was funded by Department of Science and Technology, Govt of India (CRG/2019/003037) and FRPDF scheme, Presidency University, Kolkata, India.

## Abbreviation

PTM: post translational modification
HDAC: Histone deacetylase
NAD^+^: nicotinamide adenine dinucleotide
NADP^+^: nicotinamide adenine dinucleotide phosphate
ADPr: ADP ribose
NAM: nicotinamide
(AcS): Acetyl coA synthetase
(IDH): Isocitrate dehydrogenase

---

**Table 1:** A summary of the Michaelis Menten parameters for the indicated OscobB for its deacetylase activity. Kinetic parameters of NAD^+^ as a cofactor for OscobB deacetylase activity of H3K9Ac and H3K18Ac protein from plant leaf extract as well as H3K9Ac peptide was measured using dot blot method. Reactions were carried out with 0-600 μM NAD^+^ with constant concentration of H3(60μM). Similar enzyme kinetics were calculated for acetylated mitochondrial proteins, AcS peptide and IDH2 (from extract). Data shows comparable catalytic efficiency of deacetylation in mitochondrial protein, AcS as well as in nuclear protein, H3. ± indicate S.D; n=3

| OscobB<br>Substrates | $K_m$ for NAD<sup>+</sup><br>(μM) | $k_{cat}$ (min<sup>-1</sup>) | $k_{cat}/K_m$ (M<sup>-1</sup>s<sup>-1</sup>) |
| --- | --- | --- | --- |
| H3K9Ac | 139 ± 23 | 3.0 ± 0.3 | 1.3 X 10<sup>6</sup> |
| H3K18Ac | 101 ± 33.5 | 1.1 ± 0.12 | 0.65 X 10<sup>6</sup> |
| H3K9Ac peptide | 105 ± 6 | 2 ± 0.04 | 1.14 X 10<sup>6</sup> |
| AcSAc peptide<br>(K648) | 148 ± 15.5 | 3.2 ± 0.1 | 1.3 X 10<sup>6</sup> |
| IDH2 | 191 ± 55 | 3.7 ± 0.4 | 1.16 X 10<sup>6</sup> |

## SUPPLEMENTARY FIGURES

**Fig S1.**
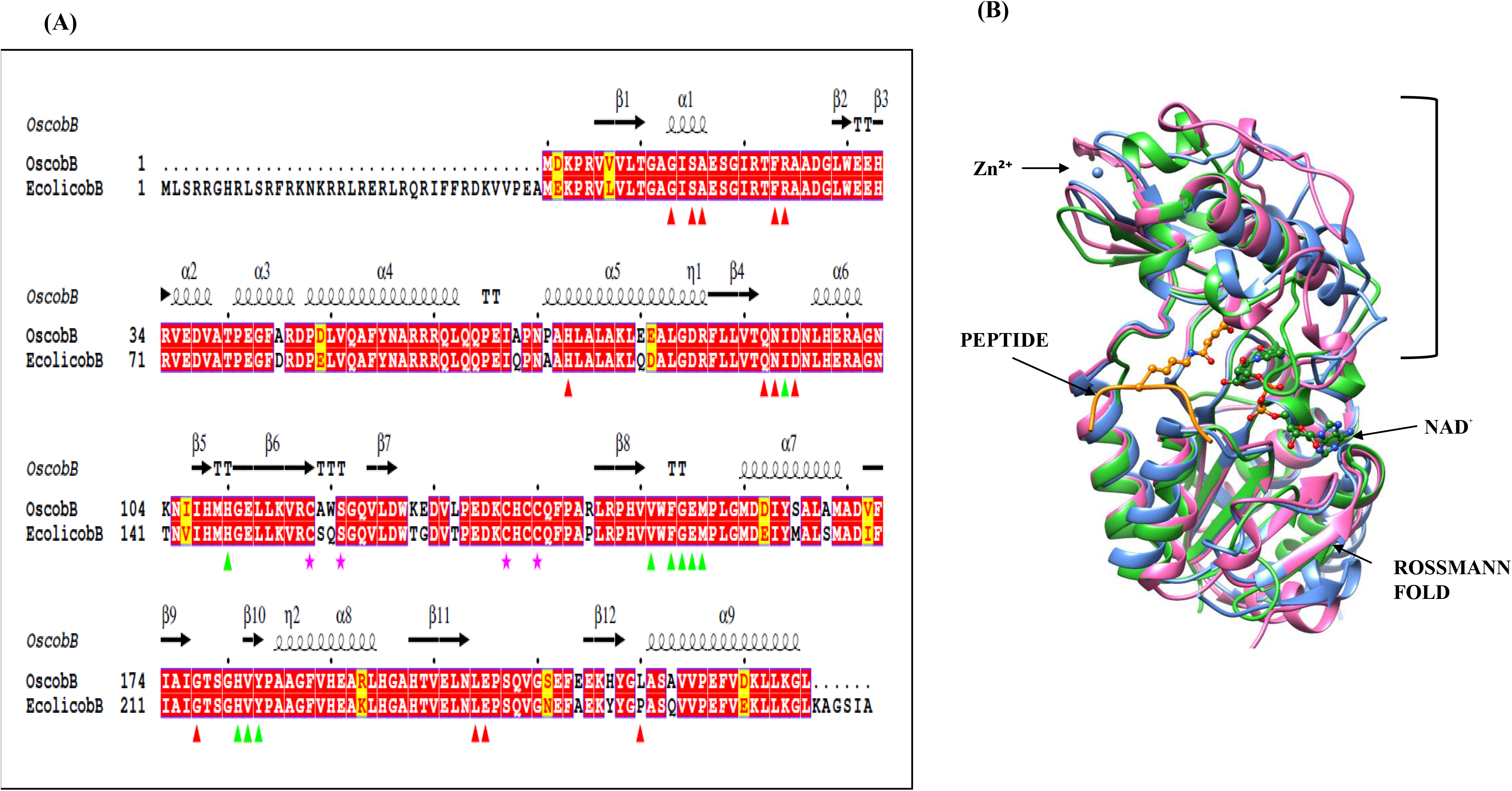
Sequence analysis of OscobB sirtuin with *E.coli* counterpart. (A) Sequence alignment of rice OscobB (Uniprot ID A2XBC4) with its homolog *E.coli* cobB (Uniprot ID P75960). The residues which line the NAD^+^ binding region are shown in red triangles, peptide binding residues with green triangles and zinc binding residues are shown with pink stars. The underlying black arrow demarcates the catalytic core region. The sequences are aligned using Clustal W and displayed using Espript 3.0. (B) Overlay of three-dimensional structures of class III sirtuin members (OscobB model, SIRT5 (3RIY) and pfSir2A (3U31)). The cartoon figure displays the Rossmann fold along with zinc ion (blue sphere), histone H3K9Ac peptide (in orange ball and stick model) and NAD^+^ (in blue ball and stick model). The Rossmann fold of all the members overlap quite well with major changes in the Zn^2+^ binding domain.

**Fig S2.**
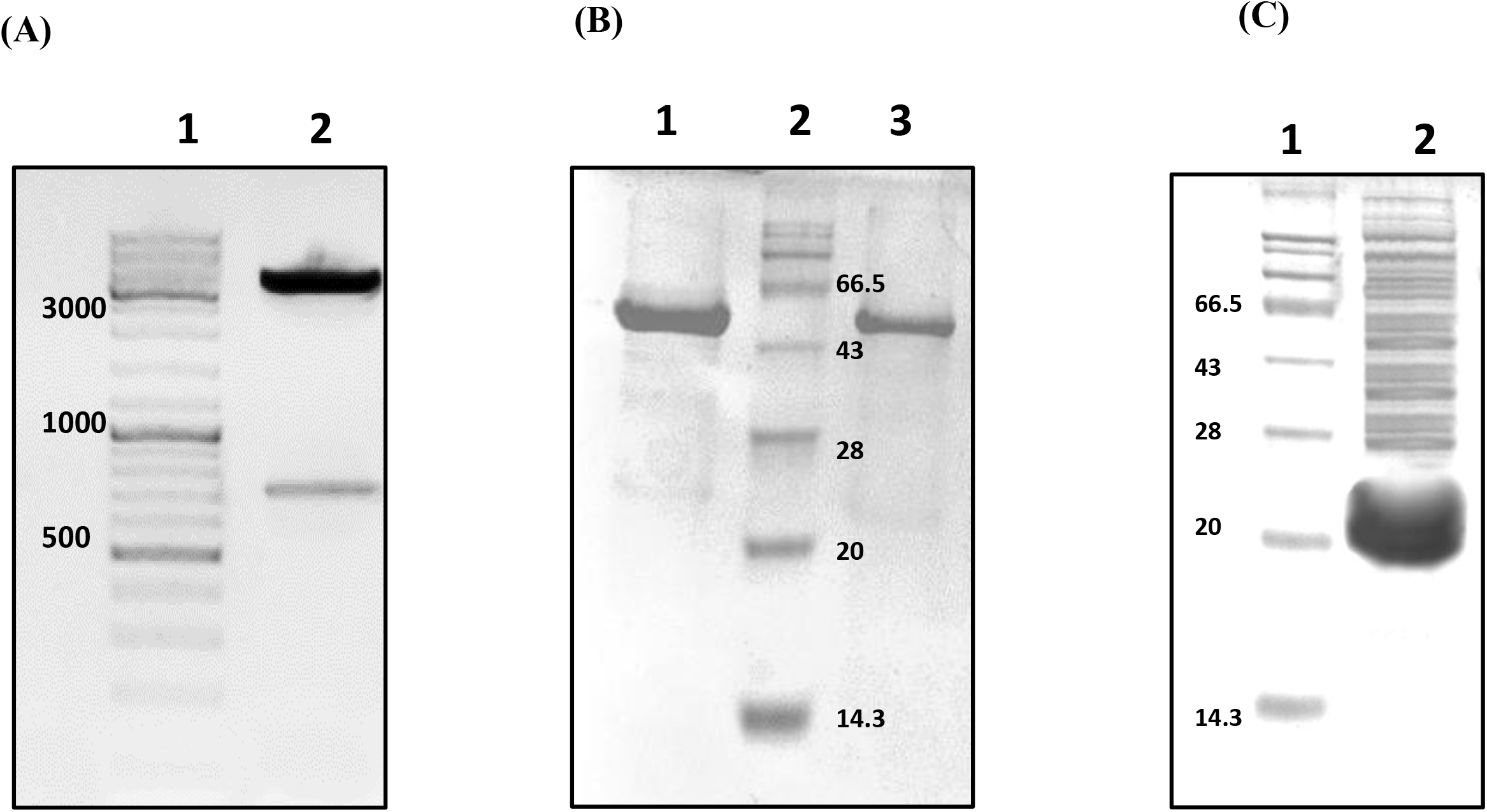
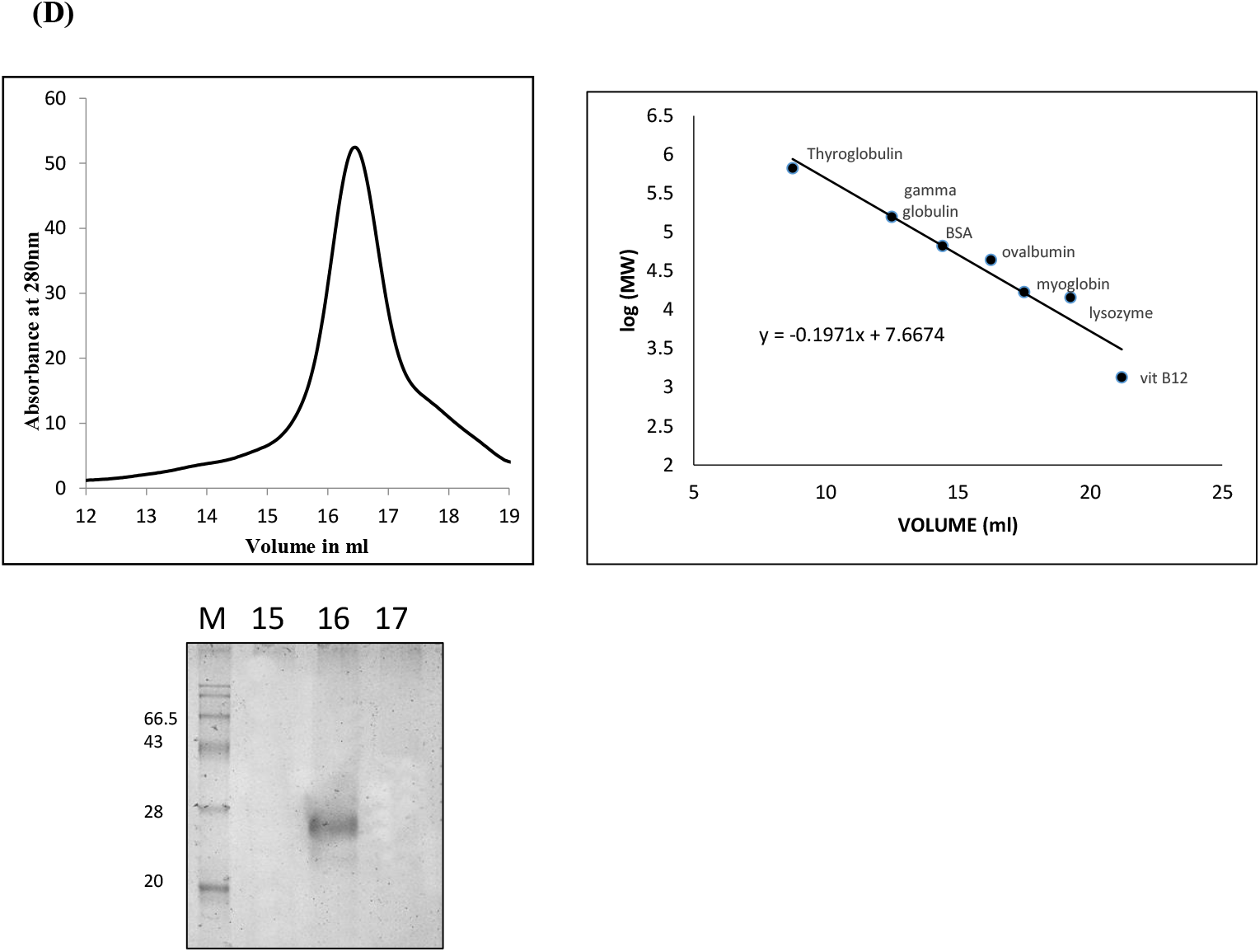
Cloning and purification of OscobB constructs. (A) 1% (w/v) agarose gel electrophoresis showing the digested product of a modified pET28a-OscobB construct. *NcoI* and *BamHI* restriction enzymes were used to digest the cloned vector. Lane 1- marker, Lane 2- double digested plasmid with OscobB insert indicating the presence of *OscobB* gene at 711bp. (B) Purification of His-tagged GST-OscobB. A coomassie blue stained 12% SDS-PAGE gel showing purified protein at 54.5kDa obtained by Nickel affinity chromatography. (C) Purification of untagged OscobB. A coomassie blue stained 12% (w/v) SDS-PAGE showing purified protein at 28kDa obtained by anion exchange chromatography. (D) Gel filtration elutes run on 12% SDS-PAGE. Lane 1 contains molecular size markers. Lanes 2, 3, 4 are elutes 15, 16 and 17 respectively where the 16^th^ elute shows SEC-purified protein suggesting that OscobB exists as a monomer. The molecular weight of OscobB was determined by a standard curve prepared from Bio-Rad protein markers.

**Fig S3.**
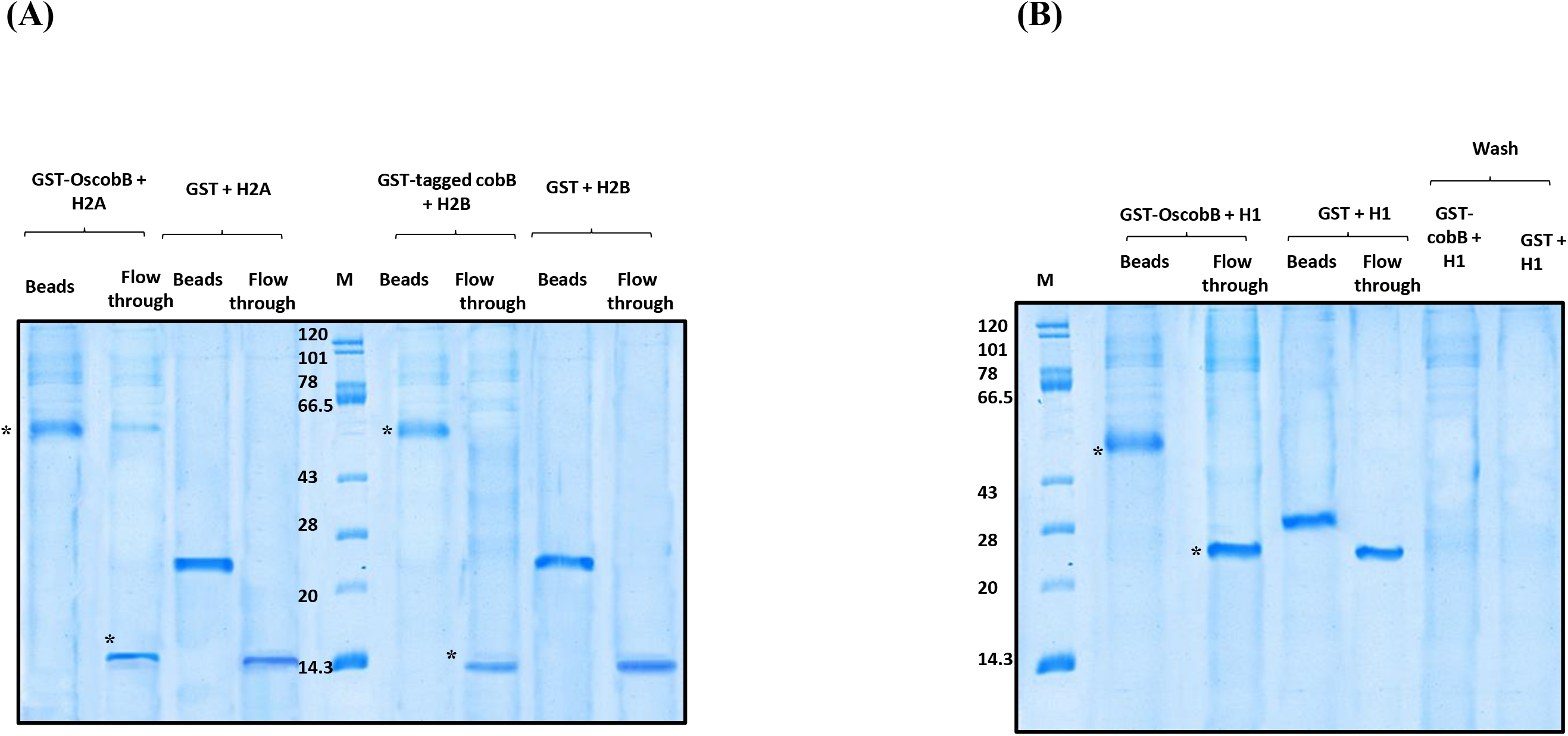
Interaction studies of OscobB with histones. (A) GST pull down binding assay. Binding studies of GST and GST-tagged OscobB with recombinant histones H2A and H2B. The histones were incubated with GST tagged proteins and allowed to be pulled down using Ni-NTA beads to extract the protein complexes. A 12% SDS-PAGE gel showed that GST-tagged OscobB was not bound to either histone, H2A and H2B and the histones were observed in the flowthrough fractions. (B) GST pull down binding assay. Binding studies of GST and GST-tagged OscobB with recombinant histone H1. Histone H1 was incubated with GST and OscobB and was allowed to be pulled down using Ni-NTA beads to extract the protein complexes. A 12% SDS-PAGE gel showed that GST-tagged OscobB did not bind to histone H1 and the histone was found in the flowthrough fraction.

**Fig S4.**
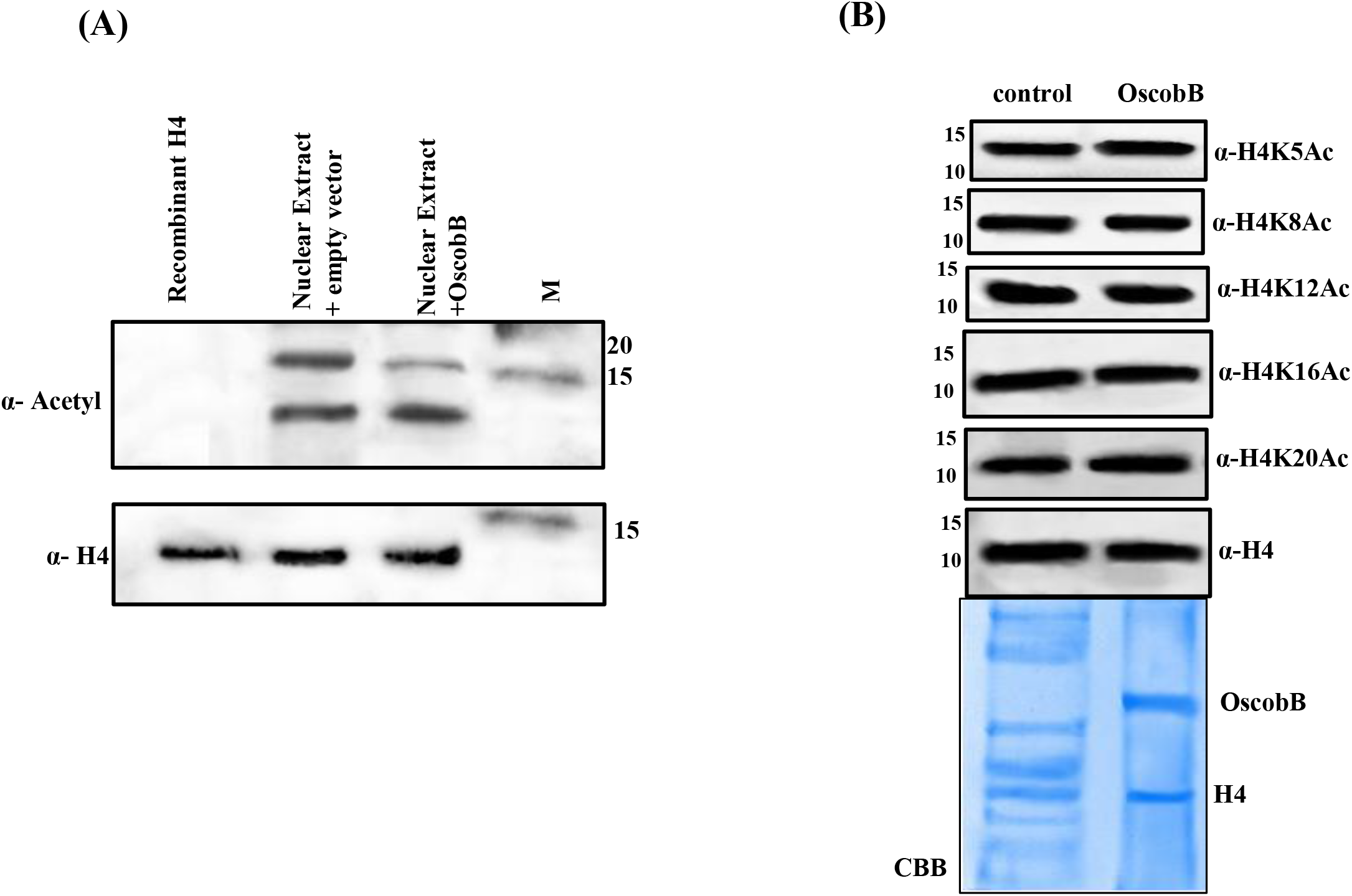
Deacetylation of histone H4 by OscobB. (A) OscobB deacetylation analysis of histone H4: The figure shows anti acetyl western blot of OscobB deacetylation reaction of nuclear extract. Here, there is no change in the acetyl band intensity of H4 in lane 3 on addition of OscobB. Empty vector in lane 2 was used as the negative control. Histone H4 is used as the loading control. (B) Detection of the deacetylation of specific lysine sites in histone H4 extracted from nuclear extract by OscobB. Lane 1- Empty modified pET28a vector, lane 2- purified OscobB. Anti-H4 antibody was used as loading control for all the western blots. A 12% SDS-PAGE gel showed purified OscobB along with histone H4 in lane 2.

**Fig S5.**
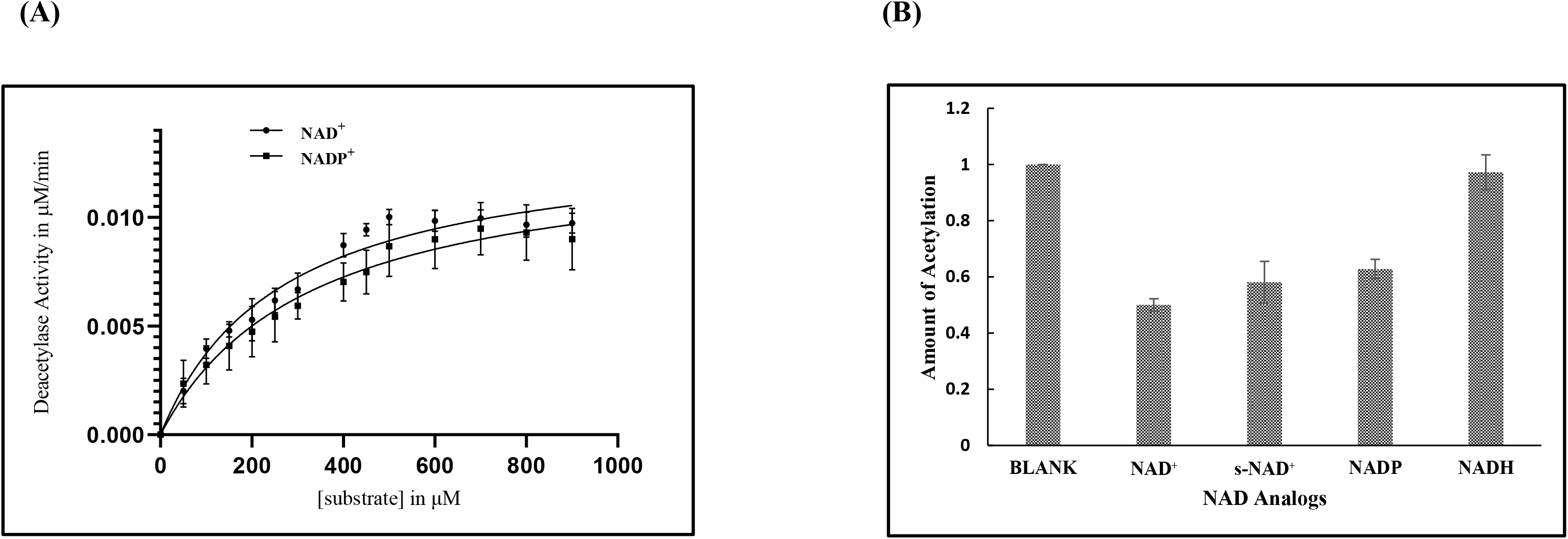
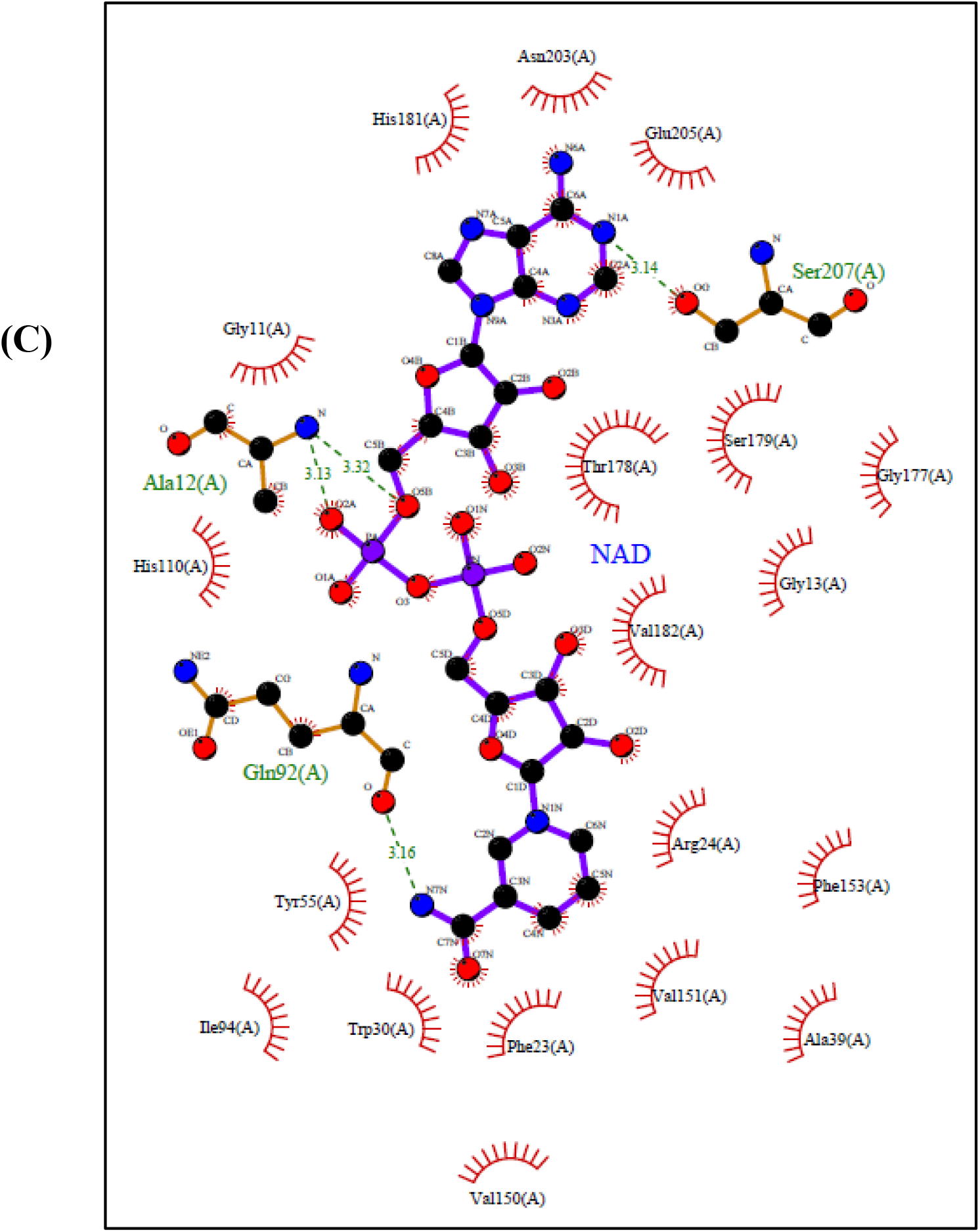
Role of NAD^+^ analogs as cofactor for substrate deacylation assay. (A) Comparative Michaelis-Menten plot of NAD^+^/NADP^+^-dependent deacetylase activity of OscobB at varying concentrations of NAD^+^ and NADP^+^ (0-800 μM) as cofactor. The reactions with histone H3 from nuclear extract were followed in triplicates using the dot blot technique. The error bar depicts the S.D.; n=3. (B) The involvement of NAD^+^ analogues as cofactor for OscobB deacetylation. The bar graph highlights the potency of the NAD^+^ analogues to act as cofactor in the deacetylation of H3K9Ac in histone H3 extracted from rice leaf nuclear extract. The reaction mixture was incubated for 2 hours and western blot analysis was done using anti-acetyl H3K9Ac primary antibody. Blank indicates the total amount of substrate. Equal amount of cofactor (450μM) was added to each reaction. (n=3; error bar: s.d.)

**Fig S6.**
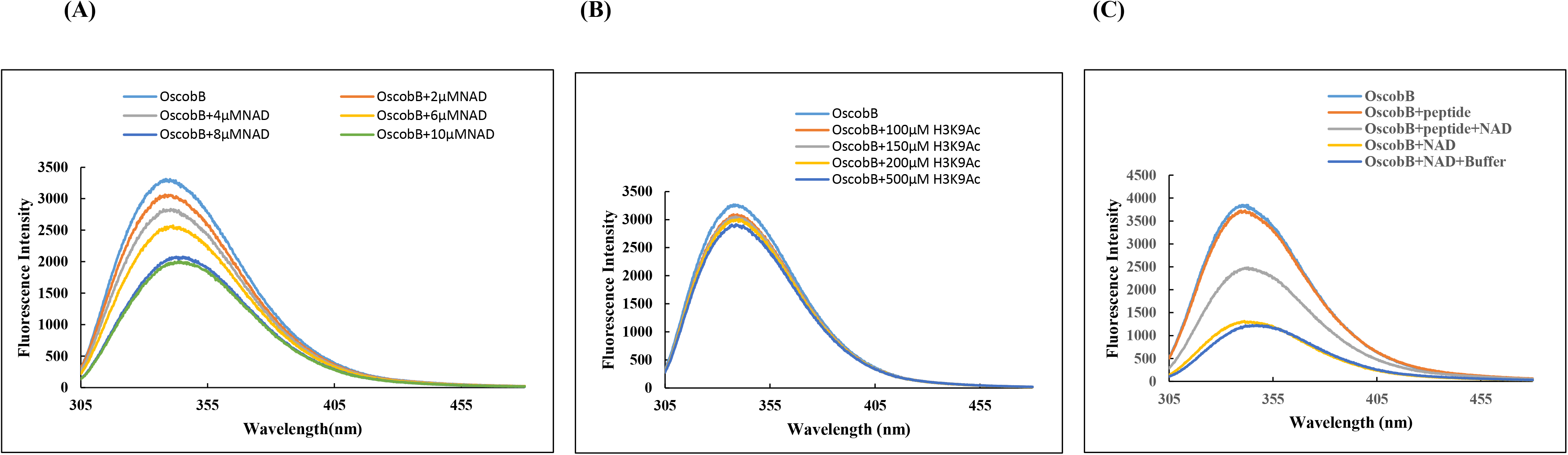
Intrinsic tryptophan fluorescence spectra of OscobB. (A) The effect of NAD^+^ on the fluorescence spectra of untagged OscobB. Increasing concentrations of NAD^+^ (0-400μM) when added to 1μM purified OscobB showed a strong binding indicated by the lowering of fluorescence intensity at 340nm. (B) The effect of substrate on the fluorescence spectra of untagged OscobB. Increasing concentrations of H3K9Ac peptide (0-100μM) when added to 1μM purified OscobB did not bind as effectively in comparison to NAD^+^ as indicated by the less fluctuation of fluorescence intensity at 340nm. (C) The effect of NAD^+^ and substrate on the fluorescence spectra of untagged OscobB. Increasing concentrations of H3K9Ac peptide (0-600μM) were added to 1μM purified OscobB and 600μM NAD^+^. The further lowering of fluorescence intensity in the presence of bound NAD^+^ indicates that binding of the cofactor might result in conformational changes in the protein resulting in stronger binding of the peptide.

**Fig S7.**
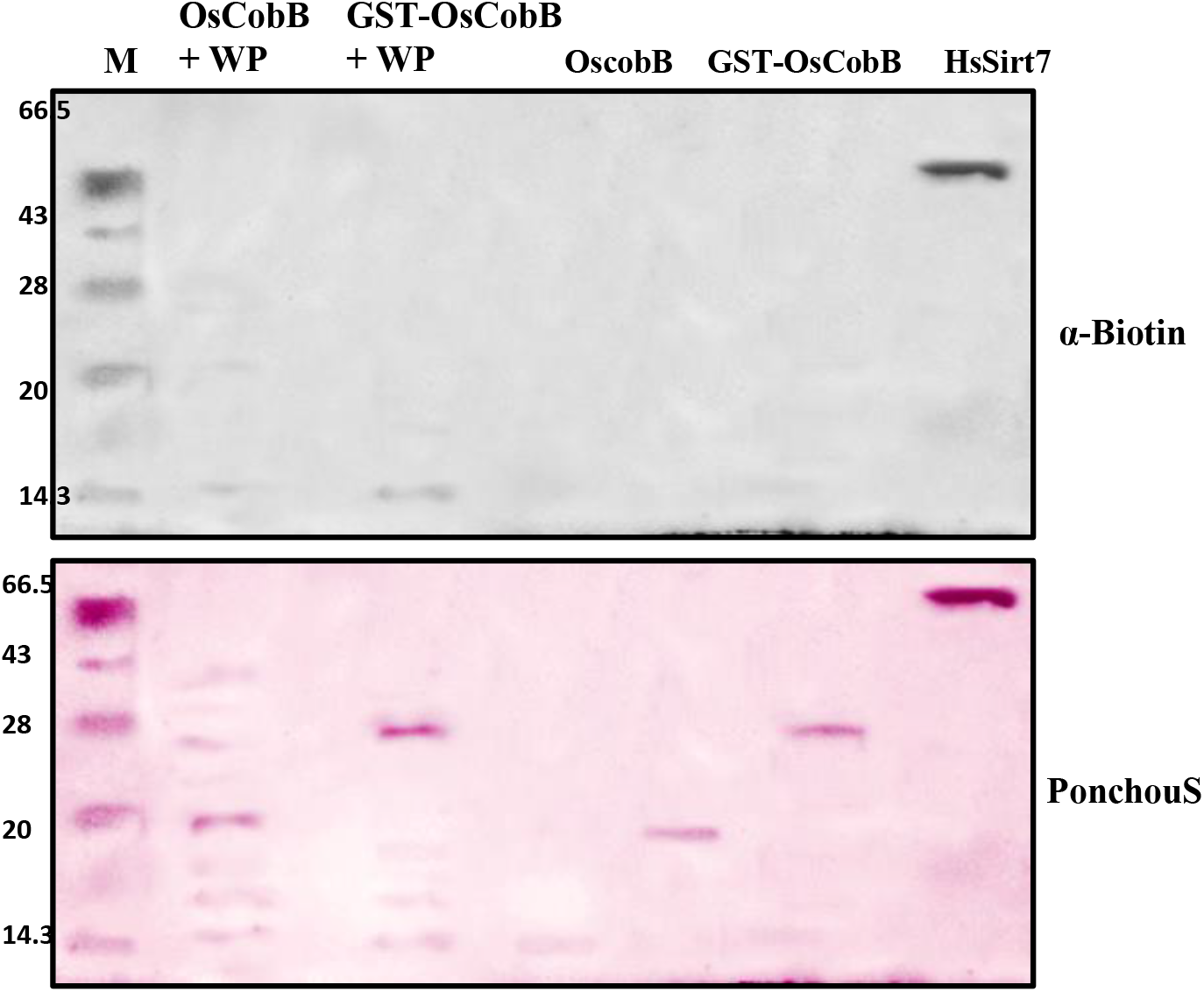
Auto-ADPr transfer reaction of the OscobB: This reaction of OscobB was carried out using Bio-NAD^+^ (40 μM) as a substrate to catch the transfer of biotinylated ADPr. The reactions were carried out with and without whole plant protein (WP). HsSIRT7 was used as the positive control. Anti-biotin western blot analysis showed the absence of ADPr transfer on any of the OscobB constructs. The reaction with whole protein from leaf extract also did not show any distinct band. The blot only showed a biotin positive band for HsSIRT7 indicating self ADP ribosylation. Ponchou S stain of the blot is indicative of the presence of purified proteins.

